# Diversity waves in collapse-driven population dynamics

**DOI:** 10.1101/022566

**Authors:** Sergei Maslov, Kim Sneppen

## Abstract

Populations of species in ecosystems are often constrained by availability of resources within their environment. In effect this means that a growth of one population, needs to be balanced by comparable reduction in populations of others. In neutral models of biodiversity all populations are assumed to change incrementally due to stochastic births and deaths of individuals. Here we propose and model another redistribution mechanism driven by abrupt and severe collapses of the entire population of a single species freeing up resources for the remaining ones. This mechanism may be relevant e.g. for communities of bacteria, with strain-specific collapses caused e.g. by invading bacteriophages, or for other ecosystems where infectious diseases play an important role.

The emergent dynamics of our system is cyclic “diversity waves” triggered by collapses of globally dominating populations. The population diversity peaks at the beginning of each wave and exponentially decreases afterwards. Species abundances are characterized by a bimodal time-aggregated distribution with the lower peak formed by populations of recently collapsed or newly introduced species, while the upper peak - species that has not yet collapsed in the current wave. In most waves both upper and lower peaks are composed of several smaller peaks. This self-organized hierarchical peak structure has a long-term memory transmitted across several waves. It gives rise to a scale-free tail of the time-aggregated population distribution with a universal exponent of 1.7. We show that diversity wave dynamics is robust with respect to variations in the rules of our model such as diffusion between multiple environments, species-specific growth and extinction rates, and bet-hedging strategies.

## Author Summary

The rate of unlimited exponential growth is traditionally used to quantify fitness of species or success of organizations in biological and economic context respectively. However, even modest population growth quickly saturates any environment. Subsequent resource redistribution between the surviving populations is assumed to be driven by incremental changes due to stochastic births and deaths of individuals. Here we propose and model another redistribution mechanism driven by sudden and severe collapses of entire populations freeing up resources for the growth of others. The emergent property of this type of dynamics are cyclic “diversity waves” each triggered by a collapse of the globally dominating population. Gradual extinctions of species within the current wave results in a scale-free time-aggregated distribution of populations of most abundant species. Our study offers insights to population dynamics of microbial communities with local collapses caused e.g. by invading bacteriophages. It also provides a simplified dynamical description of market shares of companies competing in an economic sector with frequent rate of bankruptcy.

## Introduction

Mathematical description of many processes in biology and economics or finance assumes long-term exponential growth [1, 2] yet no real-life environment allows growth to continue forever [3, 4]. In biology any growing population eventually ends ups saturating the carrying capacity of its environment determined e.g. by nutrient availability. The same is true for economies where finite pool of new customers and/or natural resources inevitably sets a limit on growth of companies. Population dynamics in saturated environments is routinely described by neutral “community drift” models [5, 6] sometimes with addition of deterministic differences in efficiency of utilizing resources [7].

Here we introduce and model the saturated-state dynamics of populations exposed to episodic random collapses. All species are assumed to share the same environment that ultimately sets the limit to their exponential growth. Examples of such systems include local populations of either a single or multiple biological species competing for the same nutrient, companies competing to increase their market shares among a limited set of customers, etc. Specific case studies can be drawn from microbial ecology where susceptible bacteria are routinely decimated by exposure to bacteriophages (see e.g. [8, 9] and references therein), or paleontological record, where entire taxonomic orders can be wiped out by sudden extinctions happening at a rate independent of order size [10].

## Model

Population growth *P*(*t*) of a single exponentially replicating species in a growth-limiting environment is traditionally described by Verhulst’s [4] or logistic equation *dP/dt* = Ω *· P ·* (1 − *P*/*P*_*tot*_), where the carrying capacity of the environment *P*_*tot*_ without loss of generality can be set to 1. In this paper we consider the collective dynamics of multiple populations competing for the same rate-limiting resource:

- Local populations are subject to episodic random collapses or extinctions. The probability of an extinction is assumed to be independent of the population size. Immediately after each collapse the freed-up resources lead to the transient exponential population growth of all remaining populations *P*_*i*_. The growth stops once the global population ∑_*j*_ *P*_*j*_ reaches the carrying capacity *P*_*tot*_ = 1.
- The environment is periodically reseeded with new species starting from the same small population size *γ* ≪ 1 (measured in units of environment’s carrying capacity).

We initially assume that the growth rates and collapse probabilities of all species are equal to each other. We also disregard the neutral drift [5] in sizes of individual populations during the time between subsequent collapses. All these assumptions will be relaxed, simulated, and discussed later in the paper (see Supplementary Materials S1 Text, S1-S7 Figures). The number of species in the steady state of the model is determined by the competition between the constant rate of introduction of new species and the overall rate of extinctions in the environment that is proportional to the number of species. To simplify our modeling we will consider a closely related variant of the model in which the number of species *N* is kept strictly constant. In this case each extinction event is immediately followed by the introduction of a brand new species. We have verified that the two version of our model have very similar steady state properties. The dynamics of the fixed-*N* model is described by

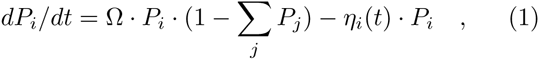

where *η*_*i*_(*t*) is the random variable which is equal to zero for surviving populations and has a large positive value for populations undergoing an extinction/collapse.

To speed up our simulations we do not continuously calculate Eq. (1) since most of the time the carrying capacity of the environment is saturated when local populations do not change. Instead we simulate the model at discrete time steps marked by extinction events. At every time step a randomly selected local population goes extinct and a brand new species with population *γ* ≪ 1 is added to the environment. We then instantaneously bring the system to its the carrying capacity by multiplying all populations by the same factor.

## Results

In spite of its simplified description of the ecosystem disregarding pairwise interactions between species our model has a rich population dynamics. Figure 1A shows the time-courses of populations in a system with *N* = 20 species and *γ* = 10^−4^. At certain times the carrying capacity of the environment is nearly exhausted by just one dominant species with *P*_*max*_ ≃ 1 visible as dark red stripes in Figure 1A. When such dominant species goes extinct a large fraction of the resources suddenly becomes available and consequently all other populations increase one and the start of another diversity wave that initially is dominated by a large number of species with similar by a large ratio 1*/*(1 − *P*_*max*_). This marks the end of population sizes. In the course of this new wave these species are eliminated one-by-one by random extinctions until only one dominant species is left standing. Its collapse ends the current and starts the new wave. In Figure 1A one can clearly distinguish about 5 such waves terminated by the extinctions of dominant species #5, 15, 6, 19, and 16 correspondingly.

**FIG. 1.**
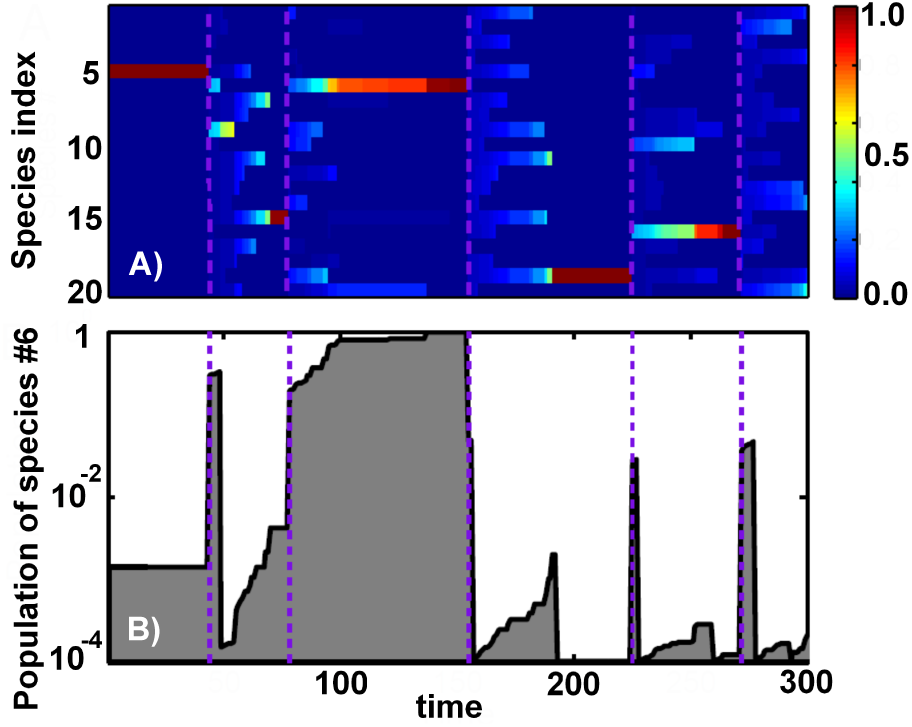
Population dynamics. The simulated model has *N* = 20 species and *γ* = 10^−4^. (A) Time-courses of populations of all species. The color denotes population sizes (see the color scale on the right) with the dominating species visible as red horizontal bands. Note five diversity waves ending at purple dashed lines. Transitions between these waves were triggered by extinctions of the dominating species # 5, 15, 6, 19, 16 correspondingly. (B) The time-course of the species # 6 with the logarithmic y-axis. Note the pattern of intermittent periods of exponential growth fueled by local extinctions.

Figure 1B shows the time-course of just one local population of the species #6 on a logarithmic scale. Between time steps 100 and 150 the population of this species nearly exhausts the carrying capacity of the environment. Its local extinction at the time step 154 ended the third diversity wave and started the fourth one. Note somewhat erratic yet distinctly exponential growth of this species happening on the slow timescale set by the frequency of extinctions. This growth should not be confused with exponential re-population of recently collapsed environments that happens much faster (a small fraction of one time step).

Figure 2 follows the population diversity (grey shaded area) defined as 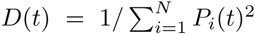 as a function of time in a system of size *N* = 1000. In general *D* can vary between ∼ 1 when one abundant species dominates the environment and *N* when all species are equally abundant. The diversity is inversely proportional to the largest population *P*_*max*_(*t*) = max_*i*_ *P*_*i*_(*t*). The diversity waves (purple dashed arrows in Figure 2) are initiated when a dominating species collapses. As a consequence, at the start of each wave the diversity abruptly increases from*∼* 1 to a substantial fraction of the maximal possible diversity *N*. After this initial burst triggered by the global redistribution of biomass, the diversity exponentially declines as exp(−*t/N*) (the dot-dashed line in Fig.2), driven by ongoing extinctions of individual populations. The typical duration, *t*_*wave*_ of a diversity wave is equal to the time required for all of the species present at the start of the wave to go extinct one-by-one. Thus it is determined by *N ·* exp(*−t*_*wave*_/*N*) *∼* 1 or

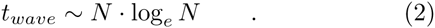

**FIG. 2.**
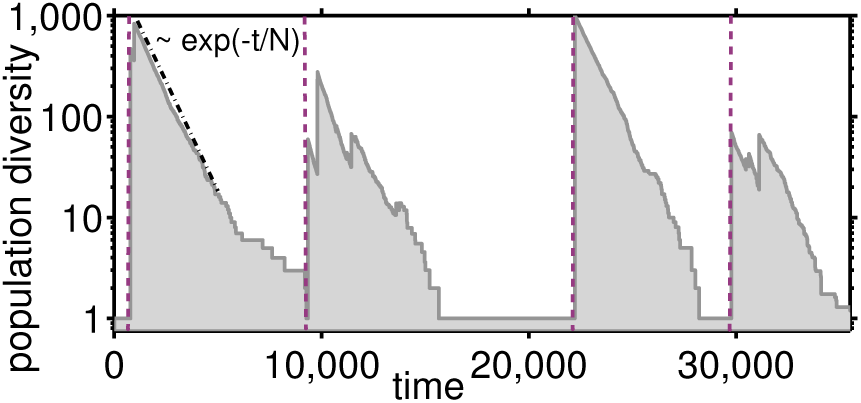
Diversity dynamics. The grey shaded area shows the the time course of the population diversity 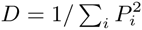 in our model with *N* = 1000 and *γ* = 10^−12^. Purple dashed lines mark the beginnings of diversity waves when a collapse of the dominant species with *P*_*max*_ ≃ 1 leads to an abrupt increase in population diversity from *∼* 1 to *∼ N*. The diversity subsequently decreases ∝ exp(*−t/N*) (dash-dotted line)

Figure 3 shows the time-aggregated distribution of population sizes for *γ* = 10^−9^ and *N* = 1000 (the grey shaded area). This logarithmically-binned distribution defined by *π*(*P*) = *d*Prob(*P*_*i*_(*t*) *> P*)*/d* log_10_ *P* were collected over 20 million individual collapses (time-steps in our model). Thus, a time-aggregated distribution is rather different from a typical “snapshot” of the system at a particular moment in time characterized by between 1 and *N* of highly abundant species in the current diversity wave. The time-aggregated distribution is bimodal with clearly separable peaks corresponding to two population subgroups. The upper peak consists of the species that have not yet been eliminated in the current wave.

**FIG. 3.**
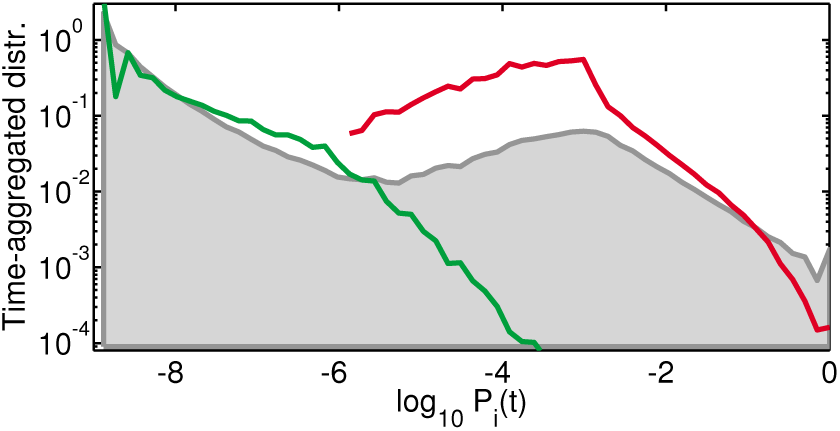
Time-aggregated population size distribution. The grey shaded area shows the time-aggregated distribution of population sizes in our model with *γ* = 10^−9^ and *N* = 1000 collected over 20 millions collapses. The green and red lines show the population size distributions collected, respectively, at the very end of each wave and at the very beginning of the next wave correspondingly as described in the text. Note that they roughly correspond to two peaks of the time-aggregated distribution.

To better understand the dynamics of the system in Figure 3 we also show the distribution of populations sizes at the very end of each diversity wave (green line) and at the beginning of the next wave (red line). That is to say, for the green curve we take a snapshot of all populations immediately after the dominant species with *P*_*max*_ *>* 1 − 1*/N* was eliminated, but before the available biomass was redistributed among all species. At those special moments, happening only once every *t*_*wave*_ time steps, most population sizes are between *γ* and *γ · N* while a small fraction reaches all the way up to ∼ 1*/N*. During the rapid growth phase immediately after our snapshot was taken, all populations grow by the same factor 1*/*(1 − *P*_*max*_) ≃ *N* thereby moving all of them to the upper peak of the time-aggregated distribution thereby starting the new wave. The red curve corresponds to the snapshot of all populations immediately after this rescaling took place. It shows that at the very beginning of the new wave local populations are broadly distributed between *∼ N · γ* and 1 with a peak around 1*/N*.

Figure 4A shows time-aggregated distributions of population sizes for *γ* = 10^−10^ and different values of *N* ranging between 100 ad 10, 000 while Figure 4B shows time-aggregated distributions with *N* = 1000 and for a wide range of *γ*. One can see that for *γ <* 0.01*/N*, the tail of the distribution for most abundant populations between 1*/N* and 1 is well fitted by a power law *π*(*P*) ∝ 1*/P*^*τ* − 1^*≃* 1*/P* ^0.7^ (dashed line in Figure 4B) corresponding to the power law distribution of population sizes on the linear scale *d*Prob(*P*_*i*_(*t*) *> P*)*/dP∼* 1*/P*^*τ*^ 1*≃/P*^1.7^. Overall Figures 4A,B demonstrate that the exponent *τ* for different values of *γ* and *N* is remarkably universal. Indeed, for a range of parameters where the upper and the lower peaks of *π*(*P*) are clearly separated, *τ* approaches a universal value *τ* = 1.7.

**FIG. 4.**
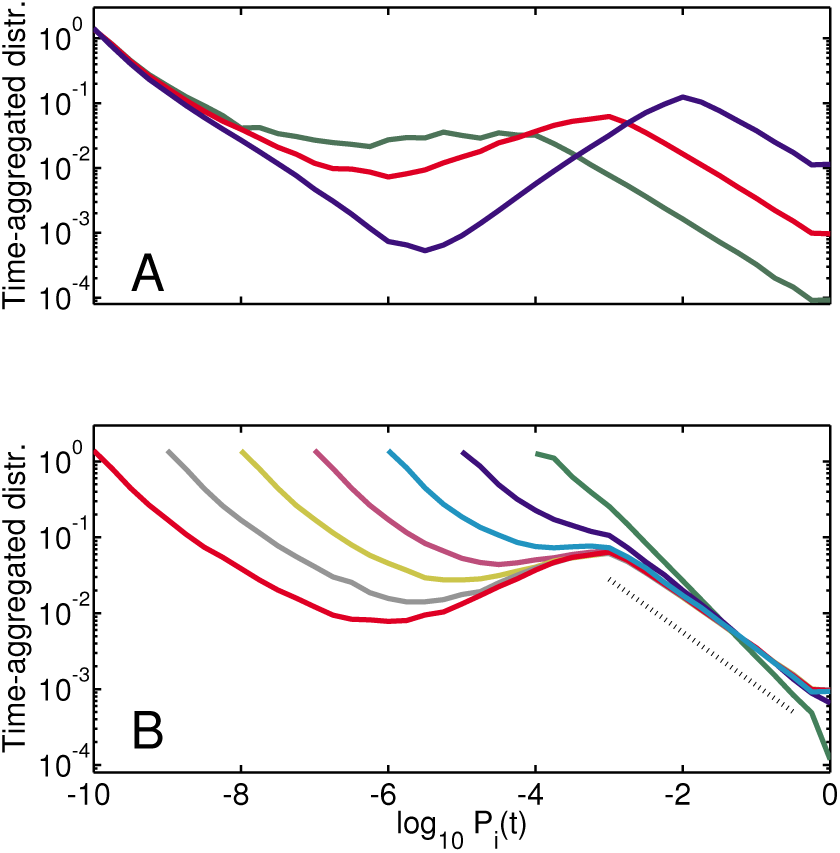
Time-aggregated distributions for different values of *N* and *γ*. Time-aggregated distributions of population sizes in our model with A) *γ* = 10^−10^ and *N* = 100 (blue), *N* = 1000 (red), and *N* = 10, 000 (green). B) *N* = 1000 and varying *γ* ranging between 10^−4^ (green) to 10^−10^ (red) in ten-fold decrements. Note the emergence of a nearly universal scale-free tail of the distribution fitted with *τ ≃* 1.7 (dashed line).

An insight into the origins of the scale-free tail of the distribution of population sizes is gained by considering a simplified version of our model in which at the start of each wave the populations of all species are artificially set to be equal to each other resulting in the maximal diversity. We further assume that *γ* ≪ 1*/N*. The passage of time *t* elapsed since the beginning of the current wave, leads to a decrease in the number of surviving species *N*_*surv*_(*t*) = *N* exp(−*t/N*), which all have the same population size *P* = 1/*N*_*surv*_(*t*) jointly filling up the carrying capacity of the environment. Above we ignore a negligible fraction (*∼ γ*) of the total population of the lower peak species. The time-aggregated probability for a species to have a population size *P*_*i*_ > *P* = 1/*N*_*surv*_(*t*) is naturally given by *N*_*surv*_(*t*)*/N* ∝ 1*/P* and thus

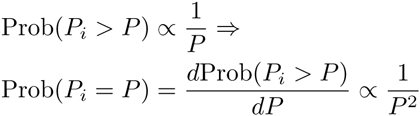

The exponent *τ* = 2 predicted by this simplified model is realized in our actual model for moderately high *γ ∼* 0.1, whereas smaller values of *γ* give rise to a different universal exponent *τ* ⋍ 1.7. The decrease of the exponent *τ* from 2 to 1.7 in our original model is the result of unequal population sizes at the beginning of a new wave. In fact, we verified numerically that *τ* = 2 is recovered if at the start of each wave one equilibrates all species abundances to 1*/N*. The first section of the S1 Text in supplementary materials provides additional details on how the reduced population diversity 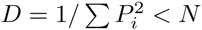 at the start of population waves affects the exponent *τ*.

Two panels in Figure 5 illustrate the difference between the simplified (panel A) and the real (panel B) models. In both versions of the model the average jump in the logarithm of surviving populations grows exponentially with time *t* elapsed since the start of the current wave: log(1 − *P*_*collapsed*_(*t*)) ⋍ *exp*(*t/N*)*/N*. However, unlike the simplified model, the population distribution in our real model has a rich hierarchical structure with multiple sub-peaks in some waves (color bands in Figure 5B). Remarkably this multi-modal distribution reappears in subsequent waves, implying that the memory about the hierarchical structure in the upper part of the distribution is transmitted to emerging populations in the lower part with sizes starting at *γ*. At the start of the next wave these same populations will move to the upper part of the distribution thereby transmitting the history across waves. Colors of symbols in Figure 5 illustrate the origin of multiple peaks. Indeed, populations in each of these peaks were born during the previous wave under similar conditions (the number of substantial populations) as described in the caption. Thus, the broadening of time-aggregated population distribution in our model compared to its history-free version is a simple manifestation of a complex interplay between “upstairs” and “downstairs” subpopulations transmitting memory across waves.

**FIG. 5.**
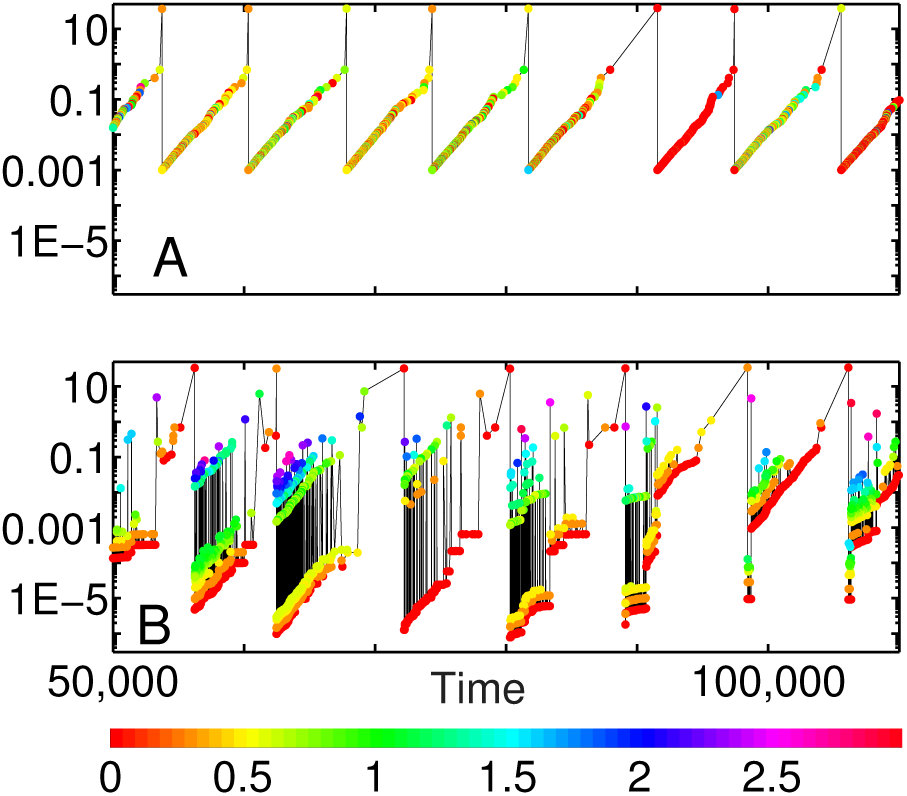
Memory of population size distribution is preserved across several diversity waves. Time course of jumps − log[1 − *P*_*collapsed*_(*t*)] in the logarithm of surviving populations following a collapse of a substantial population *P*_*collapsed*_(*t*) *>* 10^−10^ in A) the simplified model in which at the start of each wave all populations are set equal to each other; B) our basic model. Both were simulated at *N* = 1000 and *γ* = 10^−20^. Note that our basic model, unlike its simplified counterpart, preserves memory of population sizes distribution across several subsequent diversity waves. This is manifested in similar fractal structure of jumps sizes in waves #2-6 shown in panel B). Colors of symbols represent the *log*10 of the number of substantial populations during the the previous wave, when a given population originated at the small size *γ*. Thus red dots mark populations originated at the very end of the previous wave, while yellow dots those originated when there were two large populations left in the previous wave. Finally, green, blue, and purple dots refer to older populations in the previous wave.

The population distributions in both upper and lower peaks in our model are described by the same exponent *τ*. This similarity reflects the fact that individual populations in the lower peak are exposed to the same multiplicative growth as the ones in the upper peak. Finally, the intermediate region of the distribution connecting two peaks is shaped by rescaling of all populations in the lower peak as they are moved up at the beginning of a new diversity wave. When peaks are well separated (as e.g. for low values of *γ* in Figure 4) the slope of the logarithmic distribution in this region has almost exactly the same value *τ* − 1 = 0.7 and the opposite sign to the slopes in both the upper and the lower peaks.

## Discussion

In this paper we explore the population dynamics in saturated environments driven exclusively by near-complete collapses of sub-populations of competing species. This type of dynamics strongly contrasts the gradual changes implied in for example the “community drift” neutral models [5] in ecology, or for the most part incremental random walks of stock values of individual companies. Conversely, collapse-driven dynamics represents a sudden and usually large change of system composition. In ecology such collapses may be caused e.g. by invading predators or diseases, whereas in the economy, companies of any size routinely go bankrupt e.g. through excessive debt amplifying the effects of external perturbations.

First, let us consider biological systems. One of the predictions of our model is a multimodal logarithmic distribution of population sizes. Indeed, while the time-aggregated distribution is bimodal with distinct upper and lower peaks, populations within any given diversity wave cluster together in several smaller peaks persisting over several waves (color stripes in Figure 5B). This overall finding is supported by a growing body of literature [11–14] where multi-modal Species Abundance Distributions (SAD) in real ecosystems were reported for plants, birds, arthropods [14], marine organisms including single cells, corals [12], nematodes, fishes, entire seafloor communities [11], and even extinct brachiopods [15]. Like in our model, the empirical SADs range over many orders of magnitude with a noticeable depletion (or several depletions) at intermediate scales. The magnitude of this dip is usually somewhat less than predicted by our basic model but is consistent with several of its variants described below. This includes the model variant #1 inspired by the neutral theory of biodiversity [5] thought to apply to a variety of ecosystems including microbial communities [6, 7] (see S1 Figure in supplementary materials).

Needless to say, our model is not unique in generating multimodal distributions (see e.g. [13] for other examples). Conversely, some of the variants of our model give rise to interesting population dynamics including diversity waves even without any depletion in the middle of the log-binned SAD. We argue that a more reliable characterization of underlying dynamical processes can be obtained from time-series data. First, all systems capable of diversity waves are described by rapid large changes in populations of individual species. Such sudden, population-scale shifts can occur e.g. due to introduced diseases or the disappearance of keystone species [16, 17] thereby changing the composition of the entire food-web. On the microbial scale, sudden invasion of a new bacteriophage may lead to multiple orders of magnitude reduction in the population of susceptible bacteria [8, 18], potentially leading to their complete local extinction [9]. Phage-driven collapses do not spare bacteria with large populations and may even be biased towards such as postulated in the Kill-the-Winner (KtW) hypothesis [19]. The magnitude and characteristic timescales of population changes in microbial ecosystems is still being actively discussed in the literature. While Ref. [20] reports that over half of all bacterial species in marine environments varied between abundant and rare over a three-year period, other studies [21] found more modest variability at the level of species or genera over weeks to months period. However, everyone seem to agree on dramatic and rapid (often on the scale of days [22]) population shifts at the level of individual bacterial strains [8, 21, 23] caused by phage predation [22]. Except for interchangeable gene cassettes (metagenomic islands) responsible for either phage recognition cites or alternatively resistance to phages [24] these strains routinely have very similar genomes and thus may have near identical growth rates. Hence, they are capable of coexistence in the saturated state implicitly assumed in our model.

Extinctions and collapses in our model are assumed to be caused exclusively by exogenous effects such as natural catastrophes or predation by external species not sharing the carrying capacity of our environment. Real-life ecosystems can also collapse due to endogenous effects, i. e. internal interactions between species. Such intrinsic collapse mechanisms were the sole focus of earlier models by us and others (see e.g. [9, 25, 26]).

On vastly longer, geological timescales, the collapse-driven dynamics of our model resembles that of species extinctions and subsequent radiations in the paleontological record [27, 28]. One example is the recolonization by mammals of a number of ecological niches vacated by dinosaurs after the end-Cretaceous mass extinction thought to be preceded by a gradual depletion of diversity among dinosaurs who were finally wiped out by a singular catastrophic event [29]. Interestingly, the extinction rate of taxonomic orders appears to be independent of their size quantified by the number of genera they contains [10], which is also one of the assumptions of our basic model of collapse-driven dynamics.

The second area of applications of our model is to describe company dynamics in economics. The size or the market share of a publicly traded company reflected in its stock price is well approximated by a random walk with (usually) small incremental steps [30]. However, as in the case of ecosystems, this smooth and gradual drift does not account for dramatic rapid changes such as bankruptcies or market crashes. In case of companies the main driver of sudden changes is their debt [31]. When the intrinsic value of a company is much smaller than its debt, even small changes in its economical environment can make it insolvent not sparing even the biggest companies from bankruptcies (think of Enron and Lehman Brothers). Empirically, the year-to-year volatility of company’s market share varies with its size *S*, Δ*S/S ∝ S*^−0.2^ [32].

Abundance distributions in our original model and many of its variants are characterized by a power-law tail with an exponent *τ* close to 2. This is in approximate agreement with the empirical data on abundance distributions of bacteria in soil samples [33], marine viruses (phages) [34].

In the economics literature, the distributions of company sizes [35], as well as those of wealth of individuals are known to have similar scale-free tails. Recent data for companies [35] and personal wealth [36] suggest 1*/P*^1.8^ tail of the former distribution and 1*/P*^2.3^ tails of the latter one. Traditionally, scale-free tails in these distributions were explained by either stochastic multiplicative processes pushed down against the lower wall (the minimal population or company size, or welfare support for low income individuals) [37–39], by variants of rich-get-richer dynamics [40], or in terms of Self-Organized Criticality [25, 41]. The emphasis of the latter type of models on large system-wide events (avalanches [25, 41] or collapses [42]) and on separation of timescales is similar in spirit to the collapse-driven dynamics in our models. A potentially important socio-economic implication of our model is that during each wave contingent sub-peaks in the “upstairs” part of the distribution are imprinted on the “downstairs” part and thereby can be repeated in the new wave following the “revolution”.

Needless to say, our models were simplified in order to compare and contrast the collapse-driven dynamics to other mathematical descriptions of competition in saturated environments. The S1 Text in supplementary materials describes several variants of our basic model that in addition to population collapses include the following elements:

1. “Neutral drift model” assumes changes of population sizes during time intervals between collapses as described in Ref. [5]. In this model in addition to collapses a population of size *P*_*i*_ randomly drifts up and down 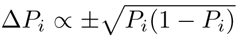. The resulting diversity waves and time-aggregated distributions can be found in the supplementary S1 Figure.
2. “Exponential fluctuations model” is another variant of the neutral scenario where the population sizes between collapses undergo slow multiplicative adjustments Δ*P*_*i*_ ∝ ±Ω_*i*_*P*_*i*_ restricted by the overall carrying capacity of the environment. Details and the resulting time-aggregated distribution can be found in the supplementary S2 Figure.
3. “Interconnected environments model” is another neutral variant of our basic rules in which spatially separated sub-populations of the same species are competing with each other for the same nutrient. Sub-populations are connected by the diffusion, that is much slower than the diffusion of shared nutrient. In this model a collapsed subpopulation is replenished by a small number *γ* of individuals diffusing from other environments, see the supplementary S3 Figure.
4. ”Kill-the-Winner (KtW) model” where collapse probability *c* systematically increase with the population size as suggested by the studies of phage-bacteria ecosystems [19]. In this particular case the diversity dynamics and the scale-free tail of the population distribution becomes sensitive to the extent that the large populations are disfavoured by collapse. When the collapse probability is proportional to population size, one obtain a flat distribution where numbers of species in each decade of population sizes are equal to each other, see the supplementary S4 Figure.
5. ”Kill-the-Looser (KtL) model”, where collapse probability *c* systematically decreases with the population size *P* as *c*(*P*) *∼ P* ^−0.2^ as suggested by the empirical studies of company dynamics [32]. As seen in the supplementary S5 Figure the diversity dynamics and the scale-free tail of the population distribution are both remarkably robust with respect to introduction of size-dependent collapse rate.
6. ”Fitness model” in which each of the species has its own growth rate Ω_*i*_ during rapid re-population phase and its own collapse probability *c*_*i*_. The supplementary S6 Figure show that the overall shape of the time-aggregated distribution is similar to that in our basic model, whereas its lower panel illustrate the interplay between population size and the the two variables that define the species’ fitness.
7. “Resilience model” as a variant of the above fitness scenario, in which collapsing species do not necessarily go into extinction. Instead, each species is assigned its own “survivor ratio” *γ*_*i*_ defined by the population drop following a collapse: *P*_*i*_ *→ γ*_*i*_*P*_*i*_. As in the previous variant each of the species is also characterized by its own growth rate Ω_*i*_. The supplementary S7 Figure shows that for intermediate populations the time-aggregated distribution is described by a power law scaling. Compared to the basic model it has a broader scaling regime and larger likelihood to have most of the “biomass” collected in one species.

Captions to supplementary S1-S7 Figures provide more detailed description of model dynamics in each of these cases. Overall, the ‘ simulations of the variants of our basic model described above preserve the general patterns of collapse-driven dynamics such as diversity wave dynamics, and a broad time-aggregated distribution of population sizes with scale-free tail for the most abundant species.

The classic definition of the fitness of a species in terms of its long-term exponential growth rate [43] is clearly inappropriate for our model. Indeed, the long-term growth rate of each of our species is exactly zero. One must keep in mind though that fitness is a very flexible term and has been defined in in a variety of ways [44] reflecting (among other things) different timescales of growth and evolution [25], and relative emphasis on population dynamics vs. risk minimization [45]. An appropriate way to quantify species’ success in a steady state system like ours is in terms of their time-averaged population size 〈*P*_*i*_(*t*)〉_*t*_.

In the last two variants of our basic model we add fitness-related parameters to each of the species: its short-term exponential growth rate Ω_*i*_(model 6 and 7), its propensity to large population collapses quantified by their frequency *c*_*i*_(model 6), and the severity of collapses quantified by surviving fraction *γ*_*i*_ of the population (model 7). Fig. 6 shows the average population size 〈*P*_*i*_(*t*)〉_*t*_ as function of Ω_*i*_ and *γ*_*i*_ in the model 6. As one can naively expects species with larger short-term growth rates and larger surviving ratios also tend to have substantially larger populations (the red area in the upper right corner of Fig. 6).

**FIG. 6.**
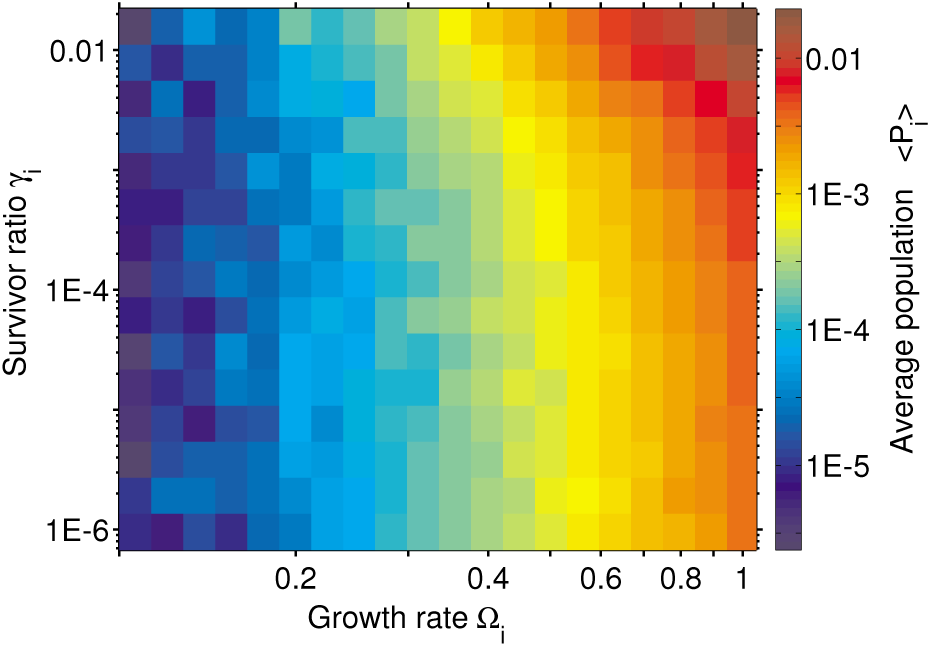
Average population vs species’ properties in the “Fitness model” variant #6. Time-averaged population of a species (see color scale on the right) plotted as a function of its re-population growth rate Ω_*i*_ (x-axis) and population drop after collapses *γ*_*i*_(y-axis). in a variant of our model with fitness differences between species. Note that the population increase with both Ω_*i*_ and *γ*_*i*_. Populations and fitness parameters of *N* = 1000 species were taken from 50 million snapshots of the model.

While in our models the probability *c*_*i*_ and severity 1/*γ*_*i*_ of collapses are assumed to be independent of growth rate Ω_*i*_ in reality they are often oppositely correlated. Indeed, in biology much as in economics/finance fast growth usually comes at the cost of fragility and exposure to downturns forcing species to optimize trade-offs.

Some environmental conditions favor the fast growth even at the cost of a higher risk of collapse, whereas other could call for sacrificing growth to minimize probability or severity of collapses. Species in frequently collapsing environments considered in our study are known to employ bet-hedging strategies [2, 45–47] to optimize their long-term growth rate. This is obtained by splitting their populations into “growth-loving” and “risk-averse” phenotypes [45, 47, 48]. One example of this type of bethedging is provided by persistor sub-populations of some bacterial species consisting of *γ ∼* 10^−4^ of the overall population [49, 50] increasing to as much as *γ* = 10^−2^ in saturated conditions (S. Semsey, private communications). A bet-hedging strategy with persistor subpopulation 10^−2^ somewhat reduces the overall growth rate (only by 1%) while dramatically reducing the severity of collapses caused by antibiotics. From Fig. 6 one infers that this is indeed a good trade-off.

In this study we presented a general modeling framework for systems driven by redistribution of rate-limiting resources freed up by episodic catastrophes. In spite of their simplicity the population dynamics in such systems happens on at least **four hierarchical timescales**. At the shortest timescale the populations grow exponentially repopulating resources vacated during a catastrophic extinction event. This exponential growth results in a steady state at which the system is poised exactly at the carrying capacity of the environment. At even longer timescales the system is described in terms of diversity waves that are the main focus of this study. These waves are an emergent dynamical property of the system in which population of surviving species grow exponentially while species diversity decays. Remarkably the information about the “upstairs”and “downstairs” population peaks survives the “revolution” at the end of each wave. This memory gives rise to the final, longest timescale in our system correlating several consecutive waves. All of this complexity is already present in our basic one-parameter model. In spite of its simplicity this model and its variants provide the foundation for future studies of collapse-driven dynamics in ecosystems, market economies, and social structures.

## SUPPLEMENTARY MATERIALS

### Fokker-Planck equation for the basic model

Let’s consider a version of our model with a very low value of *γ* to ensure the complete separation between the upper and lower peaks. The multiplicative dynamics of surviving populations in the upper peak is described by the following elementary step:

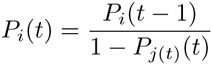

Here *P*_*j*_(*t*)(*t*) is the population at time step *t* of the species *j*(*t*) that went extinct at this time step. It can be also easily integrated for all times since the beginning of the current wave at time step 1:

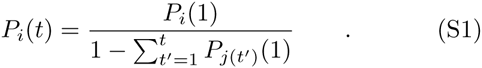

Indeed, 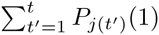 is the total initial (at time step 1) populations of all species that went extinct by the time step *t*. Hence 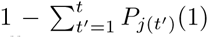 is the total initial population of all surviving species used to normalize their initial populations to give their populations at the time step *t*(population ratios of surviving species are preserved in our basic model). Taking the logarithm of both sides of Eq. S1 and approximating 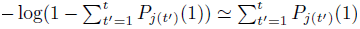, which holds as long as the system is still far away from the end of the wave 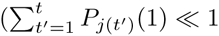, one gets:

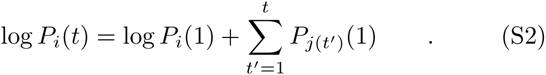

The stochastic dynamics within a single wave can thus be described by the following equation:

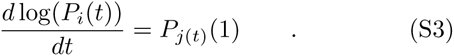

The exponent *τ* of the population size distribution in our model is determined by the balance between the noisy multiplicative population dynamics and the exponential loss of surviving species due to collapses. It can be approximated by the following Fokker-Plank-like equation:

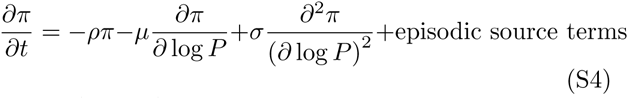

Here *π*(log *P, t*) is the time-dependent population abundance distribution in the upper peak, *ρ* the loss term due to population collapses, *μ* the logarithmic drift velocity and *σ* is the logarithmic dispersion, which is totally absent in the simplified model where all populations start at the same size.

The Eqs.S2-S3 allow us to derive the parameters of the Fokker-Plank equation in terms of the distribution of population sizes *P*_*i*_(1) at the start of the wave. Indeed, early in the wave one has *ρ* = 1*/N*, *μ* = (*P*_*i*_ (1))_*i*_ = 1*/N* and 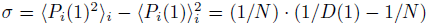 Note an unusual connection between the population diversity at the start of a wave *D*(1), and the diffusion coefficient *σ* in the Fokker-Plank equation.

The stationary solution for the time-aggregate distribution ∫ π(log *P*, *t*)*dt* has an exponential tail exp(−(*τ−*1)log *P*). It corresponds to the power law tail of the species population distribution *∝P^−τ^*. The exponent *τ* is defined by one of the two solutions to the quadratic equation

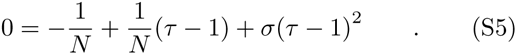

In the version of the model, where all populations at the start of the wave are equal to each other, sizes of surviving populations increase deterministically as exp(*t/N*)*/N*(see main text for the derivation) and thus have zero dispersion: *σ* = 0. Hence in this simplified version the exponent *τ* = 1 + *ρ/μ* = 2 is determined by balancing only the first two terms of this equation.

We have numerically verified that the decrease of the exponent from *τ* = 2 in the simplified model down to *τ* = 1.7 in our original model is driven entirely by noise (unequal population sizes) resulting in a finite value of *σ*. A non-zero value of *σ* in the Eq. S5 results in *τ <* 2. For example, if the populations at the start of each wave had a Poisson distribution so that *σ* = *μ* = 1*/N*, the exponent 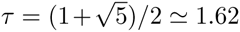 would have been defined by the solution of the golden mean equation 0 = −1+(*τ −*1) + (*τ−*1)^2^. While currently we have no first-principles argument allowing us to derive the value of *σ* in our basic model, the result from a Poisson distribution is not too far from the empirically observed exponent *τ* = 1.7

### Model variants

To test the robustness of our basic model with respect to rule changes we considered the following seven variants

1. **“Neutral drift model”**. This variant extends our basic model by adding to our standard model the random neutral drift of population sizes (Hubbell SP (2001) The unified neutral theory of biodiversity and biogeography (MPB-32), Princeton University Press) between subsequent collapse events. To simulate this random drift, at every time step the population of each species changes up or down as prescribed by dispersion of binominal distribution: 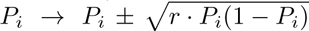, where *r* is the parameter quantifying the magnitude of fluctuations proportional to the inverse of the total population size and the square of the birth/death rate. After drift changes were applied to all populations we rescale them back to their carrying capacity ∑ *P*_*i*_ = 1. This is followed by a collapse event as in the standard model. Fig. S1 illustrates typical time courses of the diversity *D*(*t*) = 1/*∑ P*_*i*_(*t*)^2^ and time-aggregated species abundance distributions in this model variant for three values of *r* and compares them to our basic model.

**FIG. S1.**
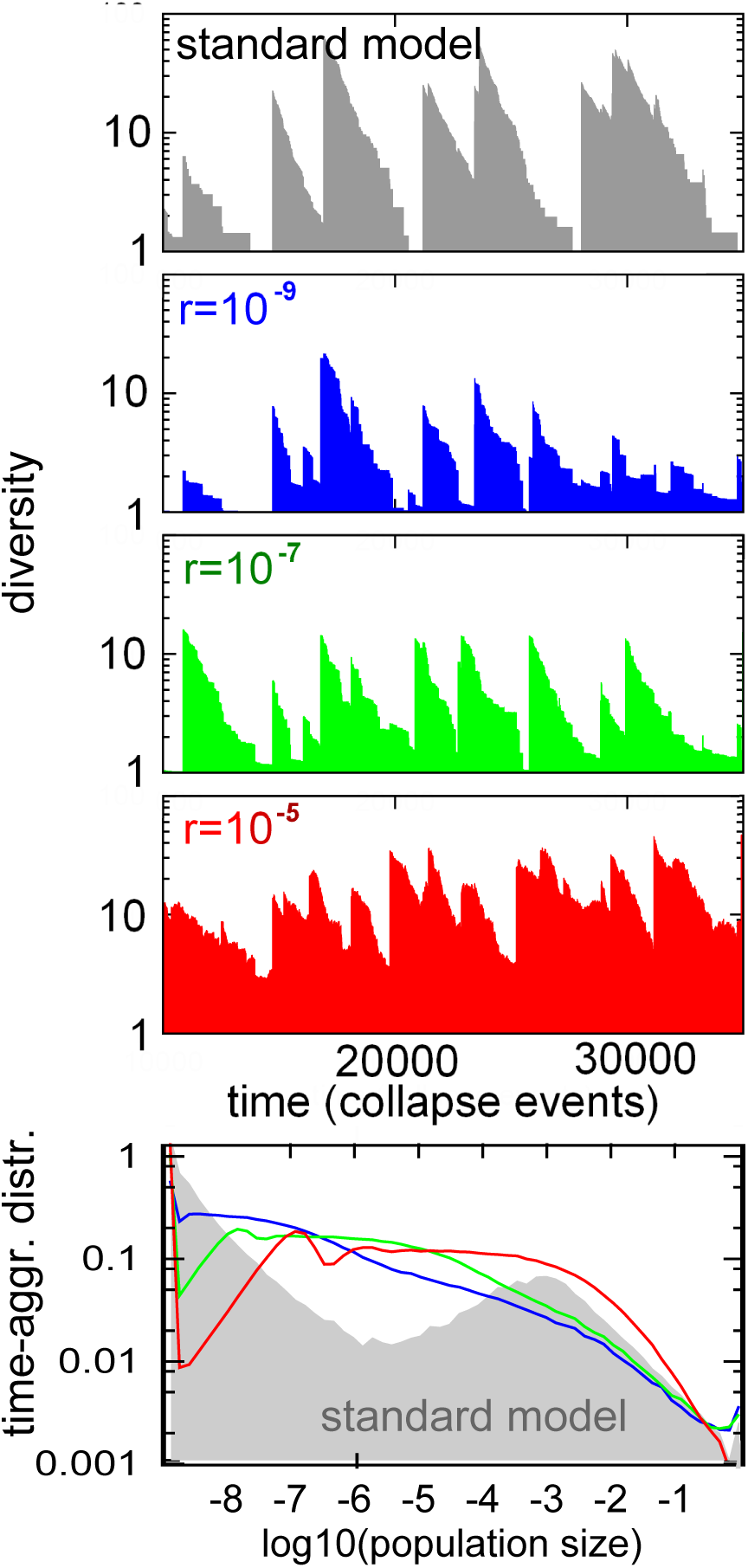
**“Neutral drift model”**. *This variant extends our basic model with N* = 1000 *and collapse ratio γ* = 10^−9^ *by adding the neutral drift at rate r taking place between subsequent collapse events in our standard model: 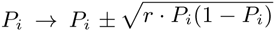. The lower panel shows the time-aggregated distributions in our system simulated for* 10^6^ *collapse events. The grey shaded area refers to our basic, unmodified model, i.e. to the r* = 0 *case, while three color lines correspond to r* = 10^−9^ *(blue), r* = 10^−7^ *(green), and r* = 10^−5^ *(red). The upper four panels illustrate typical time courses of the diversity D*(*t*) = 1*/ ∑ Pi*(*t*)^2^ *in our basic model and for three values of the r color-coded as in the lower panel.*
2. **“Exponential fluctuations model”**, where populations are exposed to random exponential shifts between successive collapse events. In this version of the model between successive collapse events all populations exponentially shift thus adjusting the way carrying capacity is divided between the. This model is similar to the “Neutral random drift” model #1 above except that changes are proportional to *P*_*i*_ and not to 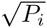. The population of each species is characterized by its own exponential rate *G*_*i*_(*t*) given by *n rand*_*i*_(*t*), where *n* quantifies the overall rate of the redistribution, while *rand*_*i*_(*t*) a random number uniformly distributed between 0 and 1 represents speciesand timespecific shifts. We assume that differences in growth rates between species are not constant but instead fluctuate on the timescale when a single species collapses. Hence, after each collapse event we reset the exponential growth we randomly reset the rates *G*_*i*_(*t*) for all species (and not just of the collapsed one). In this version of the model we also take into account the stochastic nature of the time interval *τ*_*r*_(*t*) between two successive collapse events, which is randomly chosen from the exponential distribution with mean value equal to 1. Thus between two successive collapse events each species population changes as *P*_*i*_ → *P*_*i*_ *e*^*G*_*i*_(*t*)*τr*(*t*)^ and subsequently rescaled to the carrying capacity of the environment: ∑ *P*_*i*_ = 1. As in our basic model, at every time step one randomly selected population *i* collapses and is reset to *P*_*i*_ *→ γ* while all other populations are rescaled to fill up the carrying capacity: ∑ *P*_*i*_ = 1. Fig. S2 shows the simulations of this model for several value of *n* compared to our basic (*n* = 0) model.

**FIG. S2.**
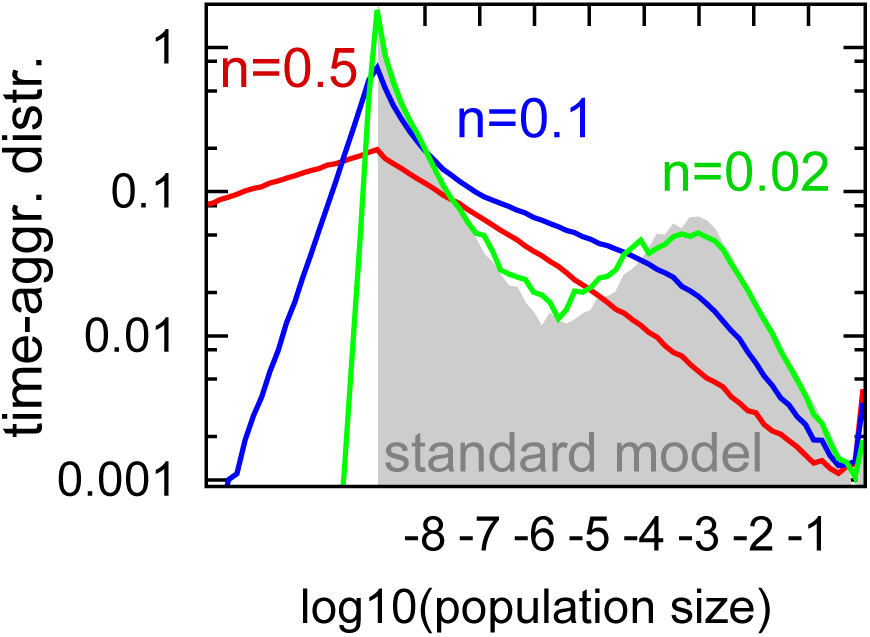
**“Exponential fluctuations model”** *with N* = 1000, *γ* = 10^−9^, and *n* = 0.02 *(green), n* = 0.1 *(blue), n* = 0.5 *(red) system simulated for* 10^6^ *collapse events. The grey shaded area shows the time-aggregated population distribution in our basic model, corresponding to the n* = 0 *limit.*
3. **“Interconnected environments model”** with diffusion. In this version of the model there is a **single** species distributed between *N* local environments. As in our basic model at every time step one local population *i* is selected for collapse and reset to *P*_*i*_ → 0 after which the populations are normalized back to the carrying capacity. The diffusion takes a fraction *γ* of the total population of 1 and distributes it equally between all local environments: *P*_*i*_ *→ P*_*i*_(1 − *γ*) + *γ/N*. Note, that here we implicitly assume that populations in all of these environments share the same carrying capacity. This is the case when diffusion rate of the rate-limiting nutrient is much faster then that of populations themselves. The main difference of this model from earlier variants is that populations in the lower peak with *P*_*i*_ ≪ 1*/N* grow approximately linearly in time (as opposed to exponentially in other versions of the model). The rate of this linear growth is the same for all species and is equal to *γ/N*. The exponential growth is restored for populations that are larger than average. Fig. S3 shows the time-aggregated distribution of local population sizes in this model variant.

**FIG. S3.**
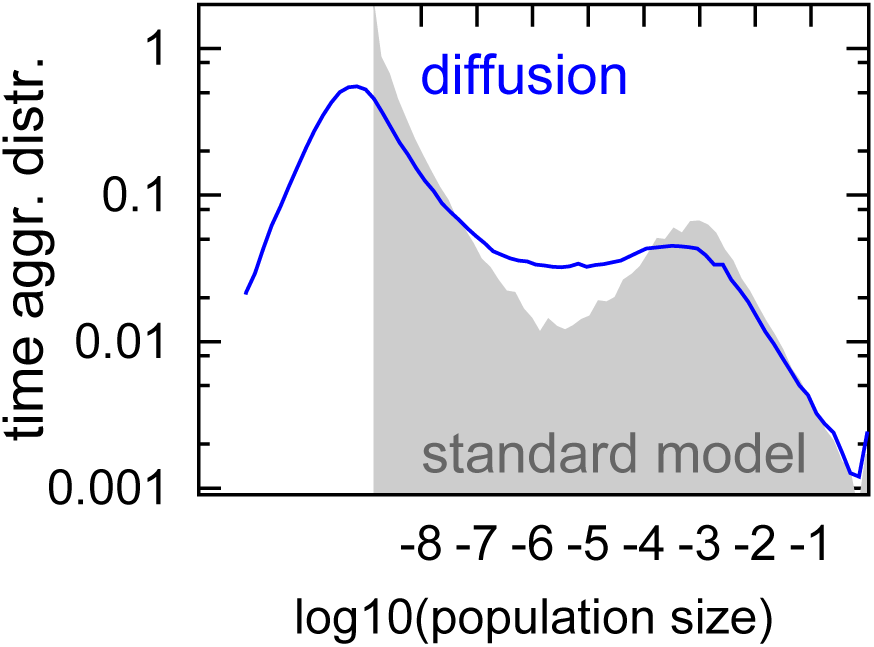
**“ Interconnected environments model”**. *Figure show time-aggregated species abundance distributions in the model with N* = 1000 *environments connected by diffusion of strength γ* = 10^−9^ *simulated over* 10^6^ *collapse events. The standard model with the same parameters is shown as the grey shaded area.*
4. **“Kill-the-Winner (KtW) model”**. For bacterial populations the direction of the trend (if any) of collapse probability with population size is currently unknown. In fact one can plausible make a case for increasing of the probability of collapse with population size due to larger populations making easier to find and overall and more attractive targets for phages. In microbiology preferential targeting of large bacterial populations by virulent phages is known as “Kill-the-Winner” (KtW) hypothesis (Thingstad TF and Lignell R (1997) Aquatic Microbial Ecology 13:19-27). Here we simulate the version of our basic model where the collapse probability systematically increases with population size. At each time step we select a random population to collapse with probability 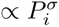. As before the collapsing population is reset to *P*_*i*_ *→ γ* and all populations are subsequently grown with equal exponential rates to complete saturation: ∑ *P*_*i*_ = 1. Fig. S4 examines time-aggregated population distributions in KtV model variant for different values of *σ*. Whereas small and moderate *σ* preserve diversity wave dynamics, the *σ* = 1 version does not exhibit diversity waves and predicts a population size distribution distributions, *dP/ds* ∝ 1*/s*, or *dP/d* log(*s*) = *constant* (equal number of species in each decade of population sizes).

**FIG. S4.**
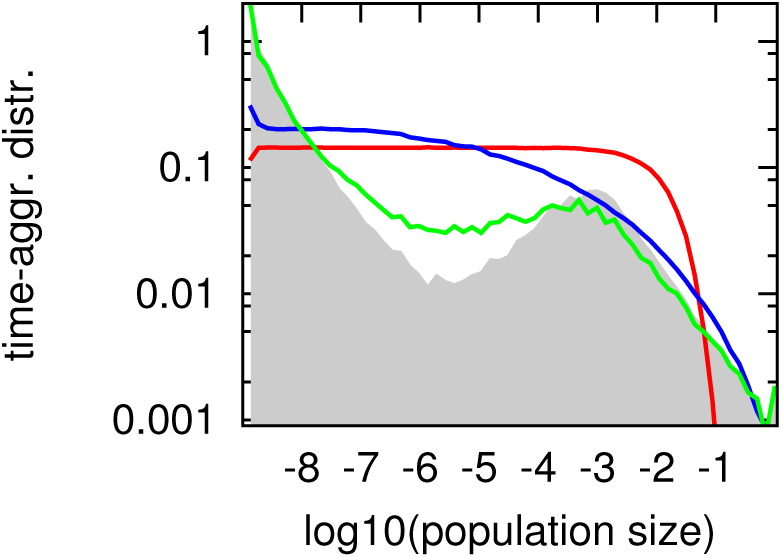
**“Kill-the-Winner (KtW) model”** *in which larger populations are preferentially targeted for collapse: 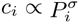. Different colors correspond to time-aggregated SADs in the model with N* = 1000, *γ* = 10^−9^, and *σ* = 0.01 *(green),* 0.2 *(blue), and σ* = 1.0 *(red) simulated over* 5·10^6^ *collapse events. The grey shaded area refers to time-aggregated population distribution in our basic, unmodified model with the same N and γ.*
5. **“Kill-the-looser (KtL) model”** in which a collapse probability declines with population size in a power-law fashion. At each time step one select a random population to collapse with probability 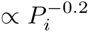 where *P*_*i*_ is the current population size of species *i*. In economics this corresponds to an intuitively plausible notion that larger companies are less likely to go bankrupt than smaller ones. Empirically, this trend is described by a power law with the exponent -0.2 (Nunes Amaral LA, Buldyrev SV, Havlin S, Leschhorn H, Maass P, Salinger MA, et al. (1997) Journal de Physique I. 7: 621-633.) Notice the emergence of the lower peak distribution distribution above *γ* that is much more narrow than in our standard model. This makes sense as in the course of each wave small populations tend to collapse over and over. These repeated collapses don’t drive other populations up by much and thus their only consequence is clustering of small populations close to the very bottom of the lower peak distribution at and above *γ*. When the dominant upper peak population finally collapses all small populations are rescaled up to form a narrow distribution around 1*/N*. This is very similar to our simplified memory-free model described in the main text and shown in Fig. 4A. The new wave starts with very high diversity *D*(*t*) ≃ *N* which is subsequently reduced with time as *D*(*t*) = *N*_*surv*_(*t*) ≃ *N* exp(−*t/N*). Here we ignore a relatively small *N*^0.2^-fold decline in collapse frequency over the range of population sizes between 1*/N* and Within the same approximation each surviving population grows as *P*(*t*) ∝ exp(*t/N*). The time-averaged distribution of populations thereby approaches the scaling regime described by:

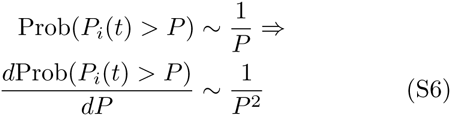 In reality the scaling exponent of the tail is around 1.8. It is the same as in our standard model but for a different reason. Indeed, taking into account that lifetime of a population before collapse scales as 1*/P*^−0.2^ = *P* ^0.2^ one gets 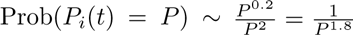

**FIG. S5.**
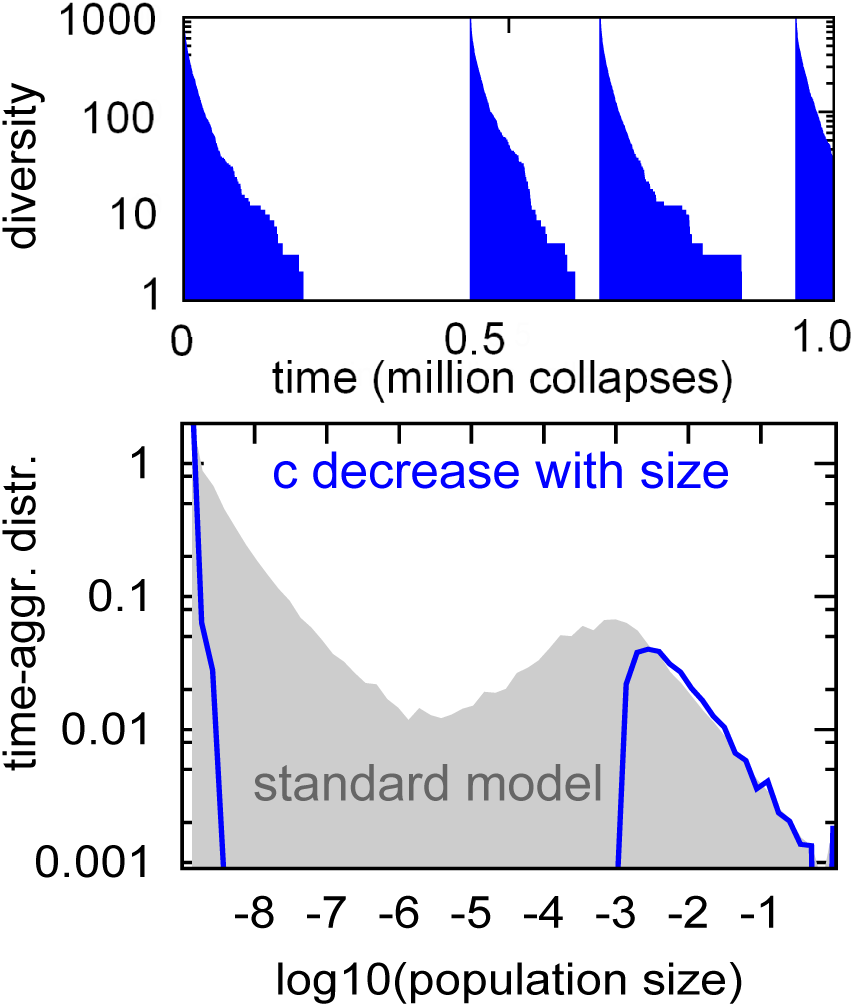
**“Kill-the-loser (KtL) model”***in which the collapse probability declines with population size. The figure shows an N* = 1000, *γ* = 10^−9^ *system simulated for* 10^6^ *collapse events. The upper panel illustrates the recurrent diversity waves, whereas the lower panel shows time-aggregated distributions, with the grey shaded area referring to our standard model.*
6. **“Fitness model”** with heterogeneous, species-specific growth rates and extinction probabilities. Each species is assigned a growth rate Ω_*i*_ used when it repopulates the freed-up carrying capacity of the environment. It also has its own extinction probability *c*_*i*_. Both Ω_*i*_ and *c*_*i*_ are logarithmically distributed in the interval between 0.1 and 1. That is to say their log10 are uniformly distributed between 1 and 0. At each time step we select one of *N* populations, with probability ∝ *c*_*k*_, this species goes extinct. It is immediately replaced by a new species with the population *P*_*k*_ *→ γ* = 10^−9^, new growth rate Ω_*i*_, and extinction probability *c*_*i*_. Sub-sequently all of the populations *i* = 1, 2*…N* are rescaled proportional to their growth rates

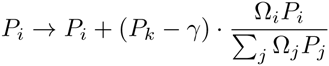

to fill up the carrying capacity of the environment ∑ *P*_*i*_= 1. The upper panel in Fig. S6 shows that the time-aggregated population distribution in this model preserves its power-law tail, whereas lower panel illustrates that in order for a species to reach substantial population size its fitness parameters need to be particularly favorable. Indeed, populations larger than 1*/N* = 0.001 tend to have smaller than average extinction probabilities *c*_*i*_, and larger than average growth rates Ω_*i*_.
7. **“Resilience model”** where heterogeneous, species-specific growth rates and survival ratios (population reduction following a collapse) are competing with each other. Each species is assigned a growth rate Ω_*i*_ ∈ [0.1, 1] and collapse size *γ*_*i*_ ∈ [10^−9^, 10^−2^], both logarithmically distributed (uniform distribution of the logarithm of the variable). At each time step we select one of *N* populations, and collapse its population *P*_*k*_ *→ γ*_*k*_ *·P*_*k*_. Note that unlike in previous versions we scale down the population proportional to its size and not proportional to the carrying capacity of the environment. This reflects a different interpretation away from our basic model, where a collapse represents not an extinction of the species followed by the appearance of a new species at a fixed (very small) population size. In the new version of the model a collapse represents a sudden but proportionate reduction of a population e.g. due to species’ phenotypic or genotypic bet-hedging. In this version of the model an extinction happens only if a very low population is reached, i.e. when *γ*_*k*_*P*_*k*_ *<* 10^−9^. If this lower bound is reached the old species goes extinct and a new species with the initial population *P*_*k*_ = 10^−9^ is introduced. The new species is assigned new random values of Ω_*k*_ and *γ*_*k*_. As in the previous model following each collapse all populations *i* = 1, 2*…N* are rescaled proportional to their growth rates 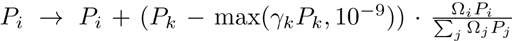 to fill up the carrying capacity of the environment:

**FIG. S6.**
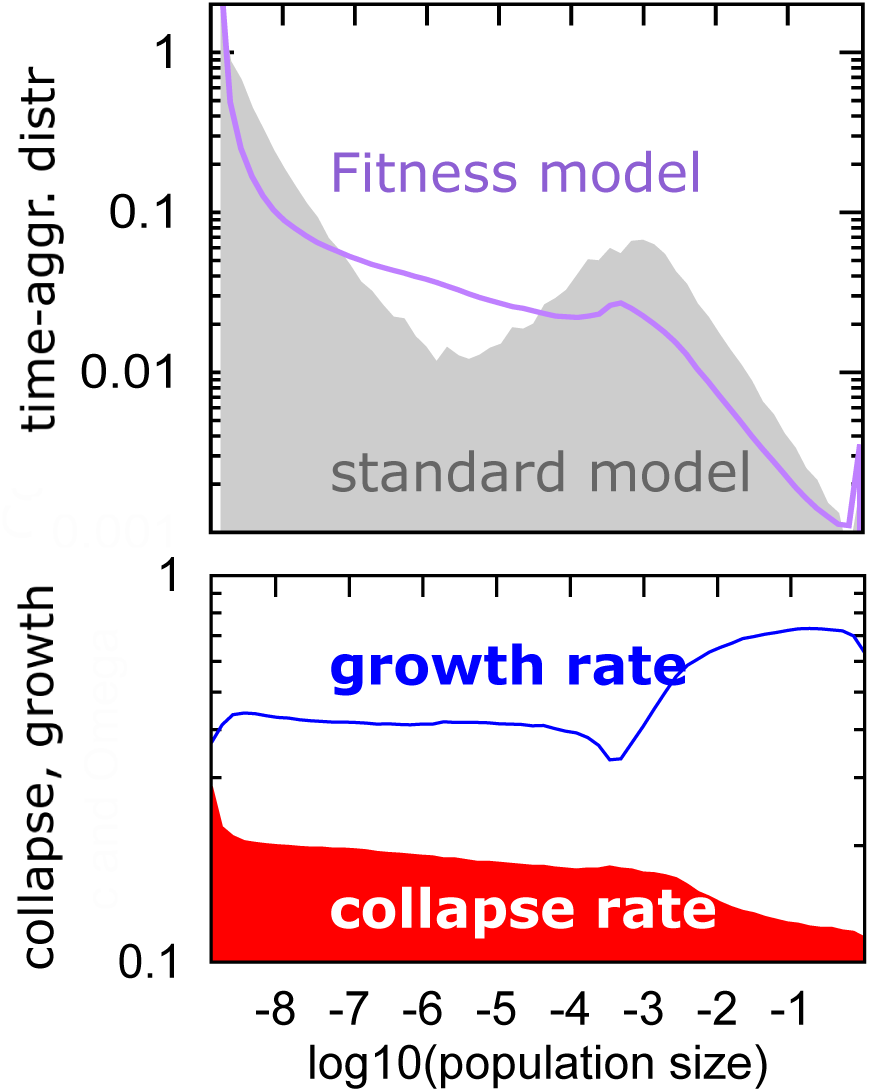
**“Fitness-model”** *with heterogeneous, species-specific growth rates and extinction probabilities. Each species is assigned a growth rate* Ω*i used when it repopulates the freed-up carrying capacity of the environment. It also has its own collapse probability ci. Both* Ω*i and ci are logarithmically distributed in the interval between 0.1 and 1. The purple curve in the upper panel shows the time-aggregated population distribution whereas the grey shaded area is that for the standard model where species’ growth and collapse rates are all equal to each other. The lower panel shows the average collapse probability and growth rate binned by the size of the population. he average growth rate* Ω*i (blue) and the average collapse probability ci (red shaded area) of species binned by their collected at every time step. Both curves represent time-aggregated averages as individual populations change with time.*

**FIG. S7.**
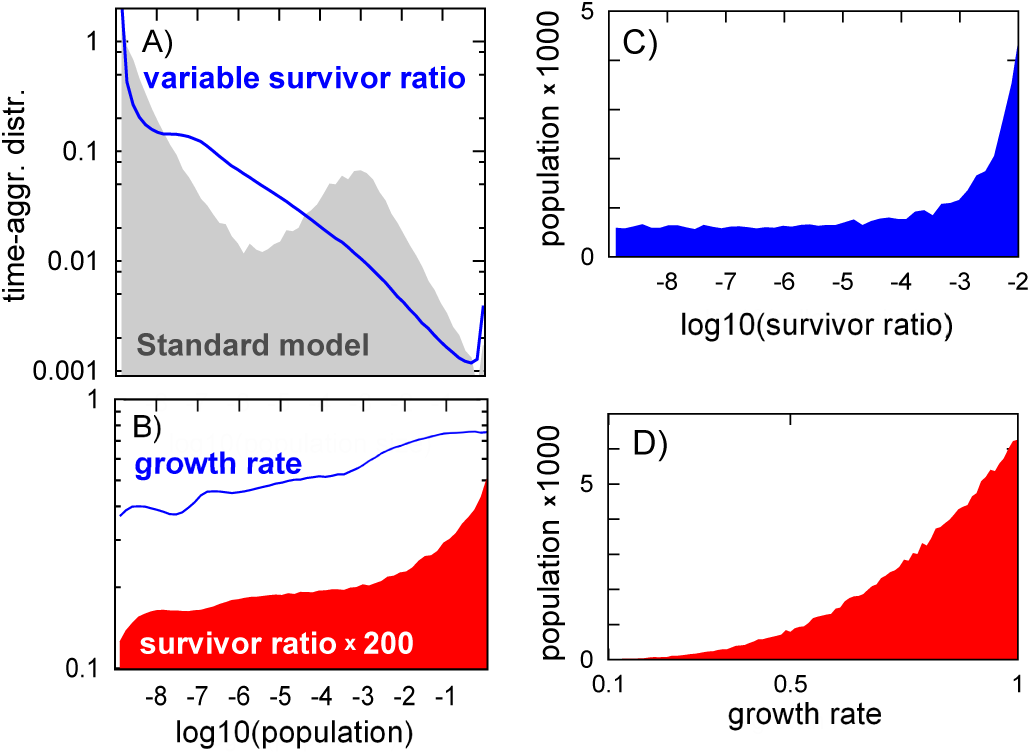
**“Resilience model”** *with heterogeneous, species-specific growth rates and survival ratios following a collapse. Each of the N* = 1000 *species is assigned a growth rate* Ω_*i*_ [0.1, 1] *and collapse size γ*_*i*_ ∈ [10^−9^, 10^−2^], *both logarithmically distributed. A) The blue curve shows the time-aggregated population distribution, whereas the grey area refers to that in our standard model from the main text. B) The average growth rate* 〈Ω〉_*i*_ *(blue) and the average survival ratio multiplied by 200 to have the same range in the plot 〈γ*_*i*_*〉 (red shaded area) as a binned by the population size collected at every time step. Both curves represent time-aggregated averages as individual populations change with time. C) The average (arithmetic) population size as a function of species’ survivor ratio Ω*_*i*_. *D) The average (arithmetic) population size as a function of species’ growth rate* Ω*i.*

